# Predicting Working Memory performance based on specific individual EEG spatiotemporal features

**DOI:** 10.1101/2022.05.06.490941

**Authors:** Vinicio Changoluisa, Claudia Poch, Pablo Campo, Francisco B. Rodriguez

**Author notes:** Corresponding author *Email addresses:* (Vinicio Changoluisa), (Francisco B. Rodriguez).

## Abstract

Working Memory (WM) is a limited capacity system for storing and processing information, which varies from subject to subject. Several works show the ability to predict the performance of WM with machine learning (ML) methods, and although good prediction results are obtained in these works, ignoring the intersubject variability and the temporal and spatial characterization in a WM task to improve the prediction in each subject. In this paper, we take advantage of the spectral properties of WM to characterize the individual differences in visual WM capacity and predict the subject’s performance. Feature selection was implemented through the selection of electrodes making use of methods to treat unbalanced classes. The results show a correlation between the accuracy achieved with an Regularized Linear Discriminant Analysis (RLDA) classifier using the power spectrum of the EEG signal and the accuracy achieved by each subject in the behavioral experiment response of a WM task with retro-cue. The proposed methodology allows identifying spatial and temporal characteristics in the WM performance in each subject. Our methodology shows that it is possible to predict the WM performance in each subject. Finally, our results showed that by knowing the spatiotemporal characteristics that predict WM performance, it is possible to customize a WM task and optimize the use of electrodes for agile processing adapted to a specific subject. Thus, we pave the way for implementing neurofeedback through a Brain-Computer Interface.

## 1. Introduction

Working memory (WM) refers to the process that allow the storage and manipulation of mental representations needed for ongoing tasks [1, 2]. It is considered to be at the base of a wide range of cognitive abilities, from perception to problem solving [3]. Despite this central role, one of the major traits of WM is its limited capacity [4, 3, 1, 5]. Whether the capacity of WM is limited by a fixed number of items or by a fixed amount of representational resource [4] is still an open debate. What it is crucial is that capacity varies among individuals[1]. Concerning individual differences, several studies have found a modulation in the activity of posterior regions as a function of the number of items held in memory [6, 7]. Another line of evidence suggests that individual differences in ignoring irrelevant information, and, as a result, restraining the number of representations held in WM, could account for differences in WM capacity [8, 3, 1]. It is widely accepted that WM performance benefits when it focuses attention on a relevant feature. One of the best known ways of orienting attention is through spatial cue [9, 10], however other features such as color, shape [11, 12, 13] or feature dimension [14, 15] can help by directing attention and modulating sensory processing areas. Traditionally, these features have been presented before the memory set (the so-called pre-cue). Limiting the number of relevant representations in WM can be achieved using a retrospective cue paradigm [10].

Different oscillatory dynamics have been associated with the performance of WM in its different phases and types of tasks [16, 17], e. g., several authors show an increase in alpha power over posterior cortical regions [18, 19, 20, 21], there is also evidence in the frontal region [22], while others mention a decrease in this alpha band [23, 24, 21] or a lateralization with an increase in alpha over posterior hemisphere ipsilateral to the direction of attention and decrease over posterior hemisphere contralateral [25, 26, 27]. Evidence shows that these alpha waves are accompanied by gamma oscillations in the prefrontal cortex [19] and in the case of alpha lateralization, it is accompanied by an increase in gamma activity in the contralateral occipital region[28]. Meanwhile, Honkanen at el. manifests presence of gamma and beta oscillations as a characteristic property of VWM maintenance at the whole cortex level [29, 30]. Similarly, several authors highlight the importance of theta an gamma in WM processes [31]. Theta band has been presented both in the posterior region, relating it to processes of attention and vision [32, 33] or in the frontal region [34, 24] relating it to WM maintenance. As can be seen, the oscillatory activity produced by WM performance is distributed by different brain regions without yet knowing the true underlying mechanisms [17, 21, 30]. Consequently, the spatiotemporal dynamics in each phase of WM performance is still not clearly understood[17, 24].

Despite not knowing exactly how brain regions are interrelated in the performance of WM, it is known that spectral characteristics show changes related to WM activity [35]. Thanks to this, it is considered as a biomarker and it has begun to be implemented in the Brain-Computer Interface(BCI) [36, 37, 38]. Taking advantage of these spectral characteristics, several works show the ability to predict the performance of WM with machine learning (ML) methods [39, 40, 41, 42, 43, 38]. Although good prediction results are obtained in these works, ignoring the inter- and intrasubject variability, there is no emphasis on feature selection, considered a key component within the classification approaches. In several of them taking all the brain regions or where the bibliography shows related activity. However, involving all electrodes from all regions is inefficient, in fact it is known that the use of a large number of electrodes does not always guarantee improved performance [44, 45, 46]. By using all the electrodes we have data with high dimensionality and noise which contains redundant information that slows down the classifier due to a positive correlation between the number of parameters to be optimized by the classifier and the number of features. The best choice of electrodes as a selection of features has been presented as an efficient alternative [47, 45, 46], but not habitually in WM tasks. Currently, ML algorithms can help a characterise and predict WM performance, and although few works have been done to date, none consider the temporal and spatial characterization in a WM task to improve the prediction in each subject. None emphasizes the need to know the moment in which each subject develops their best performance or if all subjects need a cue (e.g., precue or retro-cue), considering the spatiotemporal variability between subjects described above.

Despite the widespread use of the retro-cue paradigm, there has not been any attempt to explore which neural signals might predict behavioral performance measures [48]. We hypothesize that it is possible to predict the performance of WM in each subject and locate the spatiotemporal information in which each one will perform best, considering the intersubject variability in WM activity and the performance analysis in the retro-cue and pre-cue stimuli, which is not considered in previous works [39, 40, 41, 37, 42, 43, 38]. Our hypothesis is based on the concept that the neural activity predicts individual differences in visual WM capacity [7, 20, 43] and that the EEG spectral features are related to the WM capacity [35] therefore the differences are more clear in certain regions (and certain electrodes) in the best-performing subjects in a behavioral experiment. In the same way, we believe that not all subjects require a retro-cue and that some subjects simply use the encoding stage.

To tackle these problems, we developed a methodology to temporarily and spatially characterize a WM retro-cue task and predict performance in each subject. Using ML techniques we reveal the impact of retro-cue on each subject and the implemented methodology allowed us to locate the participating regions to achieve their best performance in each subject.

## 2. Material and Methods

### 2.1 Dataset

The dataset corresponds to the data obtained in [27], which is composed of the EEG signals of 19 adult volunteers without any history of psychiatric and neurological diseases. *Experimental Task:* The task in each trial contains four types of stages: encoding, retro-cue, probe and a black background delay interval(BBDI), shown in the upper panel in Fig. 1 (adapted from Fig.1 of [27]). A trial begins with the presentation of BBDI for 1000 ms, then in the encoding stage there are four rectangles with different orientation and color for 200 ms, these colors were randomly selected from a pool of 8 colors, without repetition in each trial. Then another BBDI stage is presented for 1000 ms, followed by a cue color (small square in the center of the screen) for 200 ms, then a BBDI is presented for 1000 ms. Finally, the probe stage presents a rectangle for 1500 ms, in which the subject responds by pressing a button if the color and orientation coincide with a rectangle presented in the encoding.

**Figure 1:**
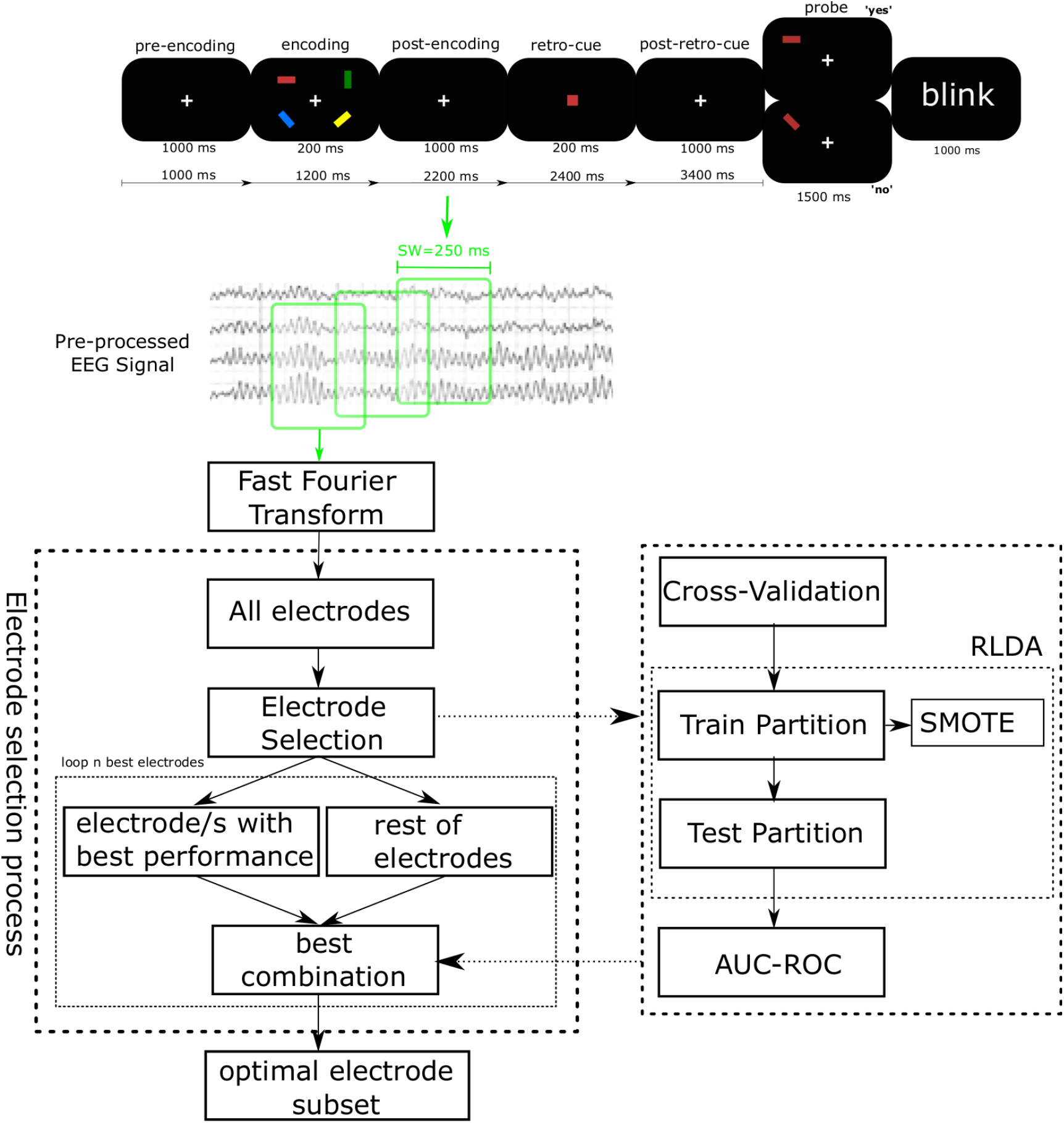
Sliding window diagram in the WM task. The upper panel shows a schematic representation of the retro-cue task (adapted from Fig.1 of [27]). The bottom panel shows the workflow of the proposed methodology.

### 2.2 EEG recording and pre-processing

Data were acquired using a Biosemi Active Two system with 128 electrodes. EOG–vertical and horizontal-electrodes and a tip-nose reference were also recorded.The data was digitalized at a sampling rate of 2048 Hz and filtered between 0.16 Hz and 100 Hz. The data were offline re-referenced to tip nose and down-sampled to 128 Hz. EEG processing was performed in MATLAB; with the FieldTrip toolbox.

After EEG recording and a first approach, the data were filtered with a bandpass filter with cut-off frequencies of 5 and 35 Hz to remove DC component and high-frequency artefacts, including power line noise (50 Hz) [49]. Simple variance-based artefact rejection is applied to reject electrodes with evident amplitude abnormalities; those were rejected with a standard deviation higher than twice the mean overall standard deviation. Thus, several electrodes’ signals were deleted for this analysis: the electrodes A24 of subject nine and B29, C5 and C30 of subject 30. Then the signal was epoched from 1000 ms before encoding onset to 2400 ms after encoding onset.

### 2.3. Classification problem: Regularized Linear Discriminant Analysis (RLDA)

The behavioral experiment was considered a binary classification problem, i.e., a class when the subject is right and another when he/she fails. Linear Discriminant Analysis (LDA) is a machine learning method [50]to deal with these types of problems. LDA is a technique that project the high-dimensional data onto a low-dimensional space to maximize class separability. To achieve optimal projection, the within-class distance is minimized and the between-class distance is maximized, simultaneously. However, classic LDA does not work well for small samples, due to the singularity of the scatter matrices involved. Regularized (RLDA) handles this problem by applying a regularization parameter (*λ*) [51]. For this, add a constant to the diagonal elements of the total scatter matrix where *λ >* 0 is known as the regularization parameter. The selection of such a parameter is crucial: small values cannot be effective in solving the singularity problem, while very large values may significantly disturb the information in the scatter matrix.

The classifier was validated using a stratified *K*-fold cross-validation scheme. The feature vector was divided into *K*-fold, one fold was used to validate and the rest (*K*-1) of folds to train. This process is repeated *K* times. The average of these operations is the accuracy reached. When stratified, it saves the proportion of data between classes. In this work *K* was equal to five.

As a performance metric, accuracy is normally used, however, it is biased for the majority classes. We use Area Under the Receiver Operating Characteristics (ROC) Curve (AUC) or (AUC-ROC curve). This metric represents the trade-off between true positive rate (TPR) and false positive rate (FPR)[52, 53] measures as follows:

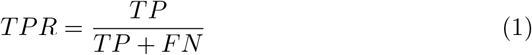

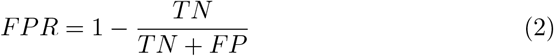

where TP, FP, TN and FN are the number of true positive, false positive, true negative and false negative, respectively. Higher the AUC-ROC, better the model is at distinguishing between the two classes.

### 2.4. Synthetic minority oversampling technique (SMOTE)

In behavioral experiments in healthy subjects, there are usually more observations of one class than another. This problem is known as class imbalance and is widely studied since they lead to two main issues: overoptimism and overfitting. Overfitting happens when the classifier is adjusted to training data and loses its ability to generalize and as a result will be unable to recognize new data.

SMOTE is one of the most used techniques to deal with this problem. The aim of this technique is to increase the number of samples of the minority class through the creation of synthetic samples. This is done by creating synthetic instances between a point of the minority class and their k-nearest minority class neighbors [54].

A frequent mistake is to apply SMOTE throughout the dataset and then divide it into testing and training set. This causes similar samples to be located in both sets, which leads to building biased models and generating overoptimistic error estimates. To avoid the aforementioned drawbacks we proceed as follows: the dataset is divided into k stratified partitions and only the training data is oversampled. In this way, it is avoided that the test samples are in the training stage.

### 2.5. Electrode selection as feature selection

Electrode selection can be understood as a feature selection technique [44, 55, 56, 45]. Among others, the main advantages of these techniques are the ability to eliminate redundant information, to reduce the number of parameters to optimize the classifier and to identify characteristics related to the targeted mental states.

Forward selection was the technique used for the feature selection. For this, we begin by evaluating the AUC-ROC of each of the electrodes, the best one is selected, then this is combined with each of the remaining ones from which the combination with the best AUC-ROC is chosen, so on until reaching the 64 best electrodes.

### 2.6. Methodology process to characterize the WM task

Sliding windows (SW) throughout the entire WM task was used. Each SW lasted 250 ms and moved every 50 ms, this sizes were arbitrarily chosen. The first window started in pre-encoding onset and the last one covered the last 250 ms before the user response onset(3400 after the pre-encoding onset). In each SW the 64 best electrodes were selected. For this, from the pre-processed EEG signal, the power spectrum of each electrode was obtained which was calculated with Fast Fourier Transform (FFT) in the frequency range 5-35 Hz. A Kaiser window was applied because it reduces spectral leakage phenomena and its good performance in EEG signals [57]. Thus, the power spectrum is the feature vector for the classifier. The next step was the selection of electrodes as explained in the previous section, successively choosing the best combination of electrodes with maximum AUC-ROC value. The workflow we follow is depicted in Fig. 1.

#### 2.6.1. Temporal representation

For the temporal analysis, we chose the two windows with a greater AUC-ROC value of each subject and grouped them into 5 groups, according to the stage: pre-encoding, encoding, post-encoding, retro-cue and post-retro-cue. In the limits of each stage, there were shared windows (which belong to two stages). These windows were assigned to the group where it has the most signal. e.g., the window starts at 900 ms and ends at 1150 ms, this window was assigned to group 2 (1000 to 1200 ms) since 100 is in group 1 and 150 in group 2.

#### 2.6.2. Spatial representation

A spatial analysis was also performed to identify the brain regions where the highest AUC-ROC values were found. For this, we create networks of the electrodes with maximum AUC-ROC obtained from a forward selection (see section 2.5). The electrodes selected have more information to predict WM activity performance and represent the networks that achieve the maximum accuracy in each subject and each group of stages according to the experiment task. We create a weighted directed graph, where each electrode represents a node linked to another node (electrode) according to the ranking achieved in the network; for example, the electrode with the best AUC-ROC will connect with the second-best this latter with the third-best, so forth. For weight, each electrode has a score, and the first electrode selected from the forward selection method has the highest ranking and continues in a descending fashion.

## 3. Results

Below we show the spatial and temporal information in which we find the maximum AUC-ROC values to identify WM with the RLDA classifier. Also, in each subject we correlated the two best AUC-ROC values achieved by classifier with the percentage of correct responses in the behavioral experiment.

### 3.1. Temporal and spatial analysis in all subjects

The top panel in Fig. 2 illustrates the temporal distribution throughout the experiment where the two highest AUC-ROC were found. In total, 38 AUC-ROCs were selected (two for each subject).

**Figure 2:**
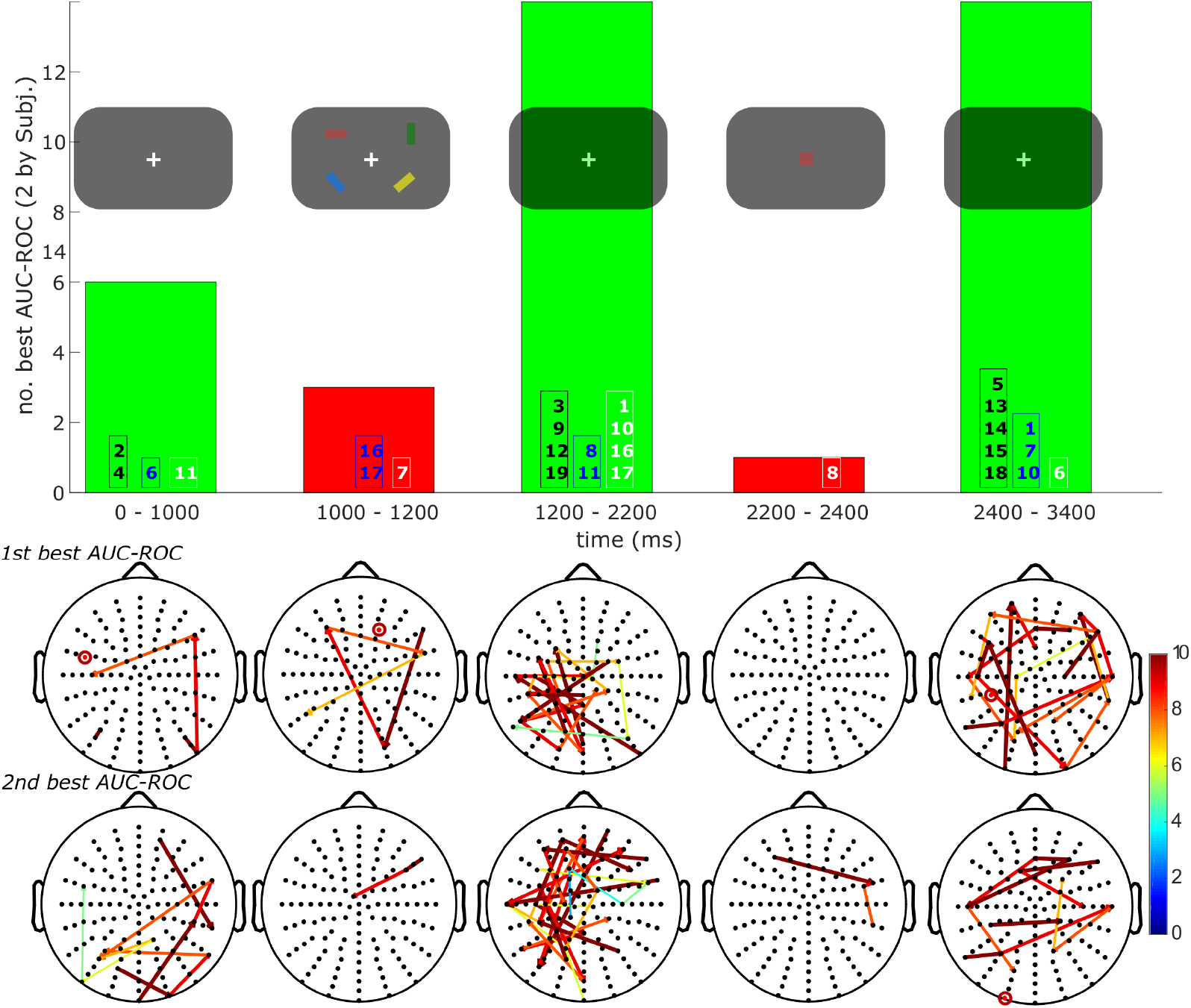
Temporal and spatial analysis of the two best AUC-ROC achieved. **Histogram (Top panel):** Each bar represents the number of best AUC-ROC values reached in each experiment stage. The numbers in the bars are the identifiers of each subject: the black color number represents when the subject has the two best AUC-ROC (1st and 2nd), blue the first best AUC-ROC and white the second. **Weighted directed graph (Bottom panel)**. Graph with the brain regions where the two best AUC-ROC values were found. Each line represents the weight of the relationship between two nodes (electrodes). The network graph represents the first link with maximum AUC-ROC with the thickest and darkest line. A circle represents an unrelated (single electrode) to achieve the best AUC-ROC.

Post-encoding and post-retro-cue are the stages that concentrate the highest amount of AUC-ROC, each with 14 values, which represents 73.6% of all selected AUC-ROC, while the encoding stage represents 7.8% and the retro-cue stage 2%. In the majority of subjects (11 of 19) the first and second best AUC-ROC coincided in the same period of time analyzed; post-retro-cue is the stage with the most matches (5 of 19).

In the spatial analysis, different activation of the electrode network were observed in each stage of the experiment. In the post-encoding stage, to achieve the first best ROC-AUC, electrode networks were activated that are mostly found in the occipital and parietal lobes. While for the post-retro-cue stage, they also involve electrodes located in the temporal and frontal lobes. Bottom panel in Fig. 2 shows the networks formed to achieve the two highest AUC-ROC values. The thickest lines and the warmest colors represent the first links created, have a weight of 10 and continue in descending order. In some cases, a single electrode was enough to achieve the highest AUC-ROC values, which are represented by a circle. On average, 3.8(*±*1.9) electrodes were necessary to achieve maximum AUC-ROC value. The second maximum AUC-ROC value was reached with 3.9(*±*2.1) electrodes.

### 3.2. Analysis of the subjects with best and worst performance in WM

According to the performance in the behavioral experiment, the dataset of 19 subjects was divided into two groups: best and worst. Taking the median as the limit between them, nine were assigned to the best performing group (on average reached 90% success) and 10 to the worst performing group (on average reached 70% success). For this analysis, the results are presented in the same way as the previous section. In the temporal analysis, it was observed that the subjects with the best performance presented a greater number of AUC-ROC values in the post-encoding stages, while those with the worst performance were presented in the post-retro-cue stage. For its part, in the spatial analysis, it maintained the pattern manifested in the previous section.

In the analysis of the best performance subjects, the post-encoding stage stands out with the greatest number AUC-ROC values with greater success (a total of eight). This stage includes the AUC-ROC value of the best performing subject in the behavioral experiment (e.g., subject 12 between others with the best performance, see Fig. 5), followed by the post-retro-cue stage, see the top panel in Fig. 3. Consider that in the pre-encoding stage, there is also a considerable number of selected AUC-ROCs, which does not happen in the worst-performing subjects, as will be seen later in Fig. 4. The bottom panel shows the brain regions where the greatest successes were found, the parietal and occipital regions are the ones that stand out the most. In this subgroup of the nine best performing subjects in WM task, to achieve maximum AUC-ROC value from the forward selection method, were necessary 4.4(*±*1.1) electrodes on average. The second maximum AUC-ROC value was reached with 3.8(*±*1.5) electrodes.

**Figure 3:**
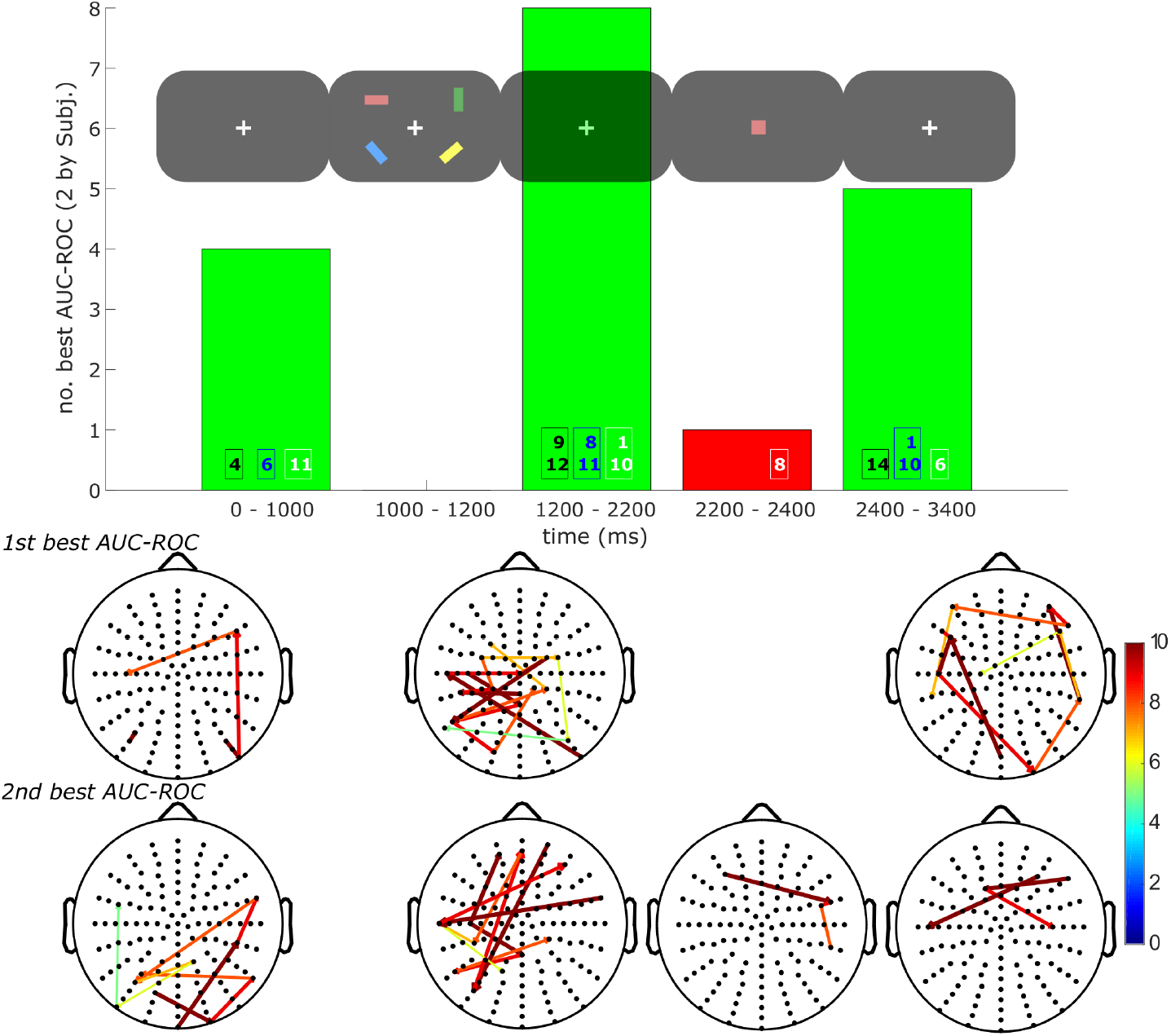
Best performing subjects data. **Histogram (Top panel):** Each bar represents the number of best AUC-ROC values reached in each experiment stage, in the same way as Fig.2. **Weighted directed graph (Bottom panel)**. Graph with the brain regions where the maximum AUC-ROC values were found for the best performing subjects.

**Figure 4:**
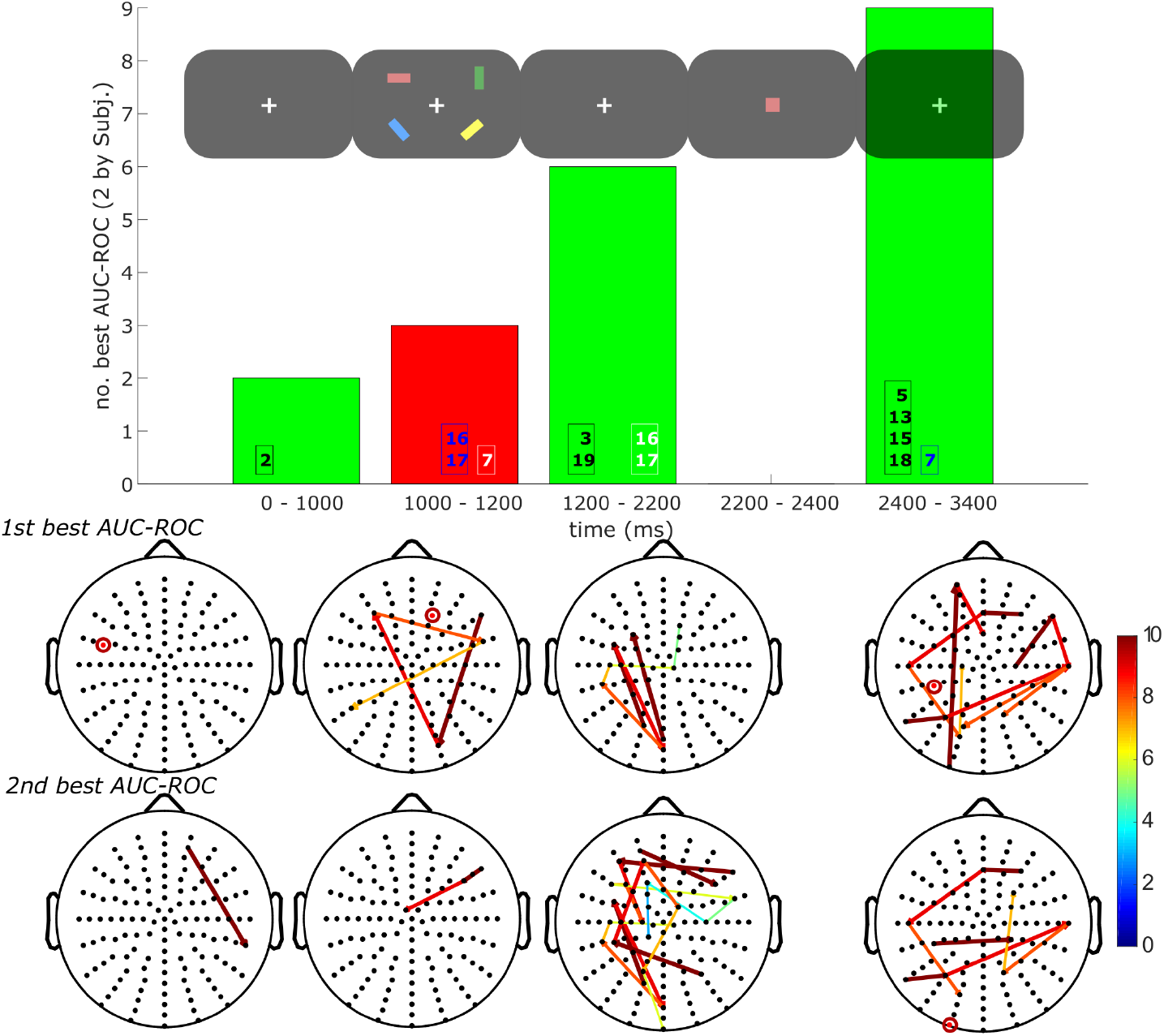
Worst performing subjects data)Histogram (Top panel): Each bar represents the number of best AUC-ROC values reached in each experiment stage, in the same way as Fig.2. **Weighted directed graph (Bottom panel)**. Graph with the brain regions where the maximum AUC-ROC values were found for the worst performing subjects.

**Figure 5:**
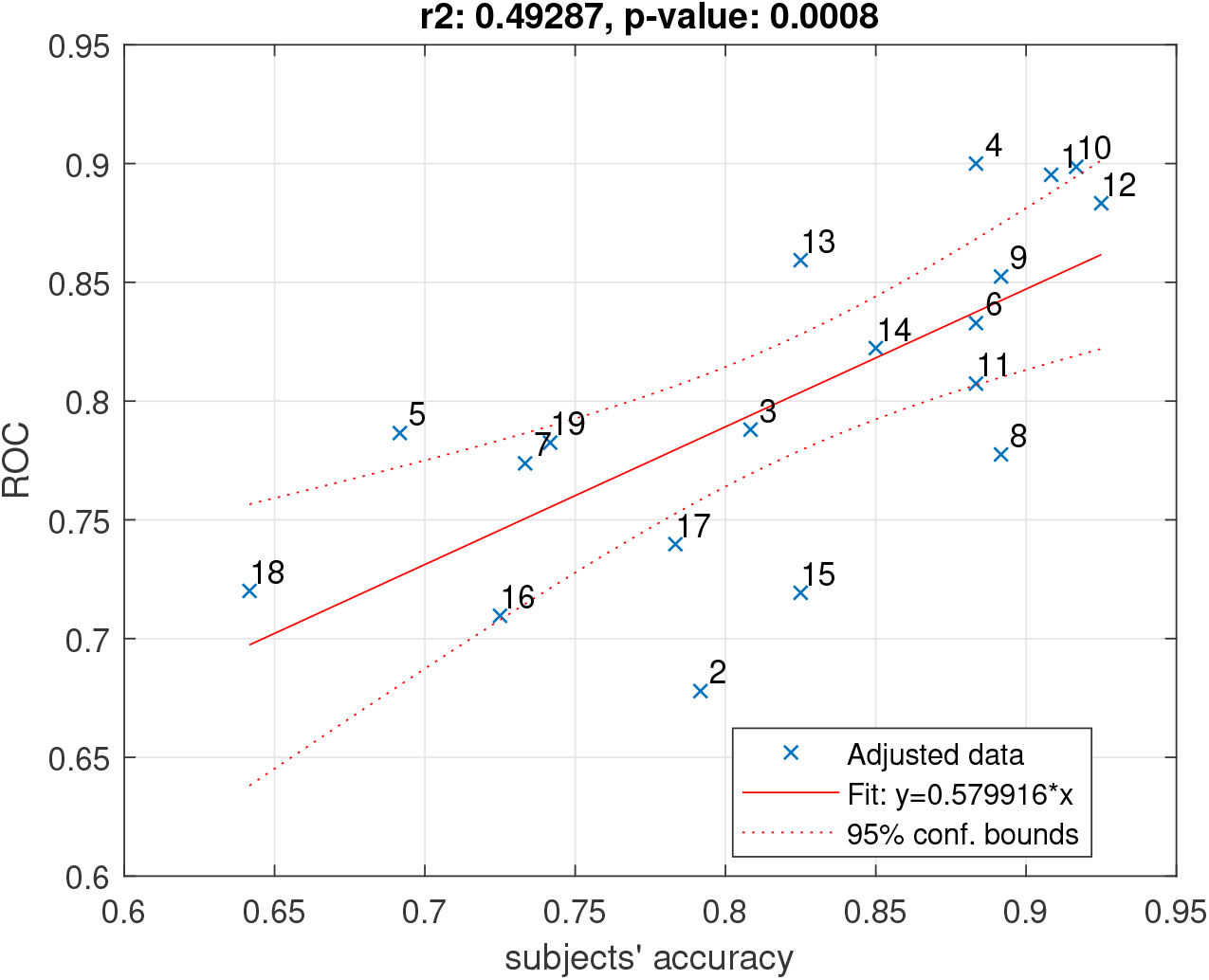
Prediction of WM performance: Linear Regression between AUC-ROC values reached by RLDA classifier and the accuracies achieved in the behavioral experiment.

On the other hand, in the analysis of the subjects with the worst performance, the post-retro-cue stage stands out with the greatest number of AUC-ROC values with greater success (a total of nine). This stage includes the AUC-ROC value of the worst-performing subject in the behavioral experiment (e.g., subject 18 or five between others with the worst WM performance, see Fig. 5), followed by the post-encoding stage, see the top panel in Fig. 4. The bottom panel highlights the brain regions with the highest number of electrodes with maximum AUC-ROC values. The occipital and parietal lobes are the ones that stand out the most, although the frontal region also participates in the post-retro-cue stage. In this subgroup of the 12 subjects with the worst WM performance, to achieve maximum AUC-ROC value were necessary, on average, 4.4(*±*2.2) electrodes. The second maximum AUC-ROC value was reached with 4.8(*±*2.8) electrodes. Although with a minimum difference, to reach the highest AUC-ROC value, there is greater dispersion in the subjects with worse performance than those with greater. To reach the second-best AUC-ROC value, it is necessary, on average, more electrodes in the subjects with worse performance than the best performance.

### 3.3. Relationship between subjects’ accuracy and classifier prediction

A linear regression was calculated to predict WM performance with EEG signals using AUC-ROC values reached by the RLDA classifier (Y) based on accuracies achieved in the behavioral experiment (X)in each subject developing a WM task. There were statistically significant differences between group means as determined by one-way ANOVA (F(1,17) = 16.52, p = 0.0008) with a R-square of 0.49. The ANOVA was performed at the confidence level of 95%, indicating that our regression model has statistically significant explanatory power. Figure 5 illustrates this relationship

## 4. Discussion

In this work we seek to predict the WM performance in each subject and analyze the spatio-temporal characteristics that are presented during a retrocue WM task. The results show a correlation between the accuracy achieved with an RLDA classifier using the power spectrum of the EEG signal and the accuracy achieved by each subject in the behavioral experiment response of a WM task with retro-cue. These findings are consistent with previous work on the use of EEG oscillations to predict WM activity [17, 16, 20, 35]. It is also supported by previous findings that relate the spectral characteristics of EEG with the WM capacity in both encoding and maintenance stages [35, 43].

The results show that it is not always necessary to have retro-cue to achieve the best performance. The temporal analysis shows that some subjects do not rely on retro-cue information. The best performing subjects in the behavioral experiment achieved their maximum AUC-ROC value in the post-encoding stage, and most did not require the retro-cue stage, while the subjects with the worse performance achieved their highest AUC-ROC value after having been presented the retro-cue. These results lead us to conclude that with adequate agile processing of the EEG signal, we can predict in real-time the subjects that require a retro-cue and those that do not, thus saving time and reducing task fatigue. Consider that in tasks that require feedback, characteristics such as strategy or speed influence WM performance [58].

For our spatial analysis, despite initially considering all electrodes from all brain regions ignoring the regions with a history of WM activity (parietal and occipital) [18, 19, 20, 21, 34, 24], our methodology found the best electrodes in these regions with evidence of WM activity. In addition, it was found that there is a contribution of the frontal region in the post-encoding stage to achieve second-best prediction. This characteristic was evidenced in all subjects, regardless of who performed better or worse in the behavioural experiment. It seems that whoever benefits from retro-cue to achieve the best performance involves a wider network of electrodes and includes, in addition to the aforementioned regions, the temporal region. The findings are consistent with the results of Griffin and Nobre [10], who mentions posterior-parietal and frontal activation during the spatial orientating of attention in retro-cue task. Also compatible with the multicomponent model proposed by Baddeley [59] and represented in a brain structure by Chai [60], in which interaction of subsystems manifests: prefrontal cortex - Central executive, the anterior cingulate cortex (ACC) - attention controller, parietal lobe - episodic buffer, occipital lobe - Visual-spatial sketch-pad. Although the true role of the frontal lobe within WM processes is still not entirely clear [61], it is unquestionable that this lobe is an active part of WM performance [61]. In recent years, several lines of evidence have challenged the assumption that PFC stores task relevant information in working memory. Instead, they propose that the role of PFC is to perform control processes necessary to the cognitive processing of the information during a working memory task [62, 63]. This control is accomplished by the interaction between PFC and posterior regions through a selective top-down transmission of information [61]. Likewise, it is important to mention that we did not analyze the frequency bands that best predict the WM performance. Despite this, our findings are consistent with previous works in which the interaction between the posterior parietal and frontal region stands out [64, 35, 65, 66]. Future works can analyze the selected electrodes and determine what other characteristics allow them to achieve the best precision and how their different combinations determine a high prediction. The results of this work show that a machine learning technique, with total ignorance of the brain regions involved in WM performance, can determine which regions most influence to achieve the best prediction of WM performance. This can be explained thanks to the extraction of characteristics used in this work, through power spectrum that has shown its robustness in the extraction of spectral patterns in EEG signals [67]. In addition to the selection of electrodes, which is understood as a feature selection technique, which has been satisfactorily tested and widely used in BCI [45, 46]. Among the main advantages of the selection of electrodes are the ability to eliminate redundant information, to reduce the number of parameters to optimize the classifier, and to identify characteristics related to the targeted mental states.

The results of this work show the need to create adaptive systems that facilitate personalized stimulation according to the needs of each individual. Elucidating the underlying mechanisms of WM will help to accurately identify spaciotemporal characteristics, eliminate redundant information, and lower computational cost. This can lead to real-time EEG monitoring devices like BCIs allowing a promising technique for neurofeedback training [68].

## Acknowledgements

This work was founded by Predoctoral Research Grants 2015-AR2Q9086 through SENESCYT, Universidad Politécnica Salesiana 041-02-2021-04-16, 034-02-2022-03-31, and Spanish projects AEI/FEDER PID2020-114867RB-I00, PID2019-111335GA-I00 (http://www.mineco.gob.es/).

